# Paracingulate sulcus morphology and hallucinations in clinical and non-clinical groups

**DOI:** 10.1101/284752

**Authors:** Jane R. Garrison, Charles Fernyhough, Simon McCarthy-Jones, Jon S. Simons, Iris E.C. Sommer

## Abstract

Hallucinations are a characteristic symptom of psychotic mental health conditions that are also experienced by many individuals without a clinical diagnosis. Research has linked the experience of hallucinations in schizophrenia to differences in the length of the paracingulate sulcus (PCS), a structure in the medial prefrontal cortex of the brain which has previously been associated with the ability to differentiate perceived and imagined information. We investigated whether this notion of a specific morphological basis for hallucinations in the paracingulate cortex extends to individuals without a clinical diagnosis by testing the hypothesis that non-clinical individuals with hallucinations have shorter PCS than non-clinical individuals without hallucinations. Structural MRI scans were examined from three demographically matched groups of individuals: 50 patients with psychotic diagnoses who experienced auditory verbal hallucinations, 50 non-clinical individuals with auditory verbal hallucinations, and 50 healthy control subjects with no life-time history of hallucinations. Measurements of paracingulate sulcal length were compared between the groups and the results verified using automated data-driven gyrification analyses. Patients with hallucinations had shorter PCS than both healthy controls and non-clinical individuals with hallucinations, with no difference between non-clinical individuals with hallucinations and healthy controls. These findings suggest that the association of shorter PCS length with hallucinations is specific to patients with a psychotic disorder. This presents challenges for continuum models of psychosis and suggests possible differences in the mechanisms underlying hallucinations in clinical and non-clinical groups.

## Introduction

Hallucinations are a common and debilitating symptom associated with several mental health disorders, but are also experienced by many individuals without a clinical disorder. Questions remain as to whether the mechanisms underlying hallucinations in clinical and non-clinical groups are the same – is there a simple continuum in the experience of hallucinations, with those with a clinical diagnosis lying at one extreme, or are there more fundamental differences in the underlying neural processes that explain the differences in perceptual experience and phenomenology between these two groups?

Hallucinations in patients with schizophrenia are often associated with impairment in reality monitoring, the cognitive ability to distinguish between real and imagined information^1,2^. With neuroimaging studies of reality monitoring in healthy individuals repeatedly revealing activity within the anterior medial prefrontal cortex (mPFC; ^3–5^), recent reality monitoring research has focused on the paracingulate sulcus (PCS), a structure that lies in the dorsal anterior cingulate region of the mPFC (Figure 1). Among the last sulci to develop *in utero*, the PCS shows significant inter-individual variation, being completely absent in 12% to 27% of healthy individuals^6–8^. People with no discernable PCS in either brain hemisphere show reduced reality monitoring accuracy compared to individuals with a visible PCS in one or both hemispheres of the brain^9^. Furthermore, an investigation of PCS morphology in patients with schizophrenia who were distinguished by whether they experienced hallucinations revealed that hallucinations were associated with a significant reduction in PCS length^10^. This suggests a specific morphological basis for these experiences within the PCS, that might be associated with an impairment in reality monitoring contributing to the attribution failure underlying hallucinations.

**Figure 1.**
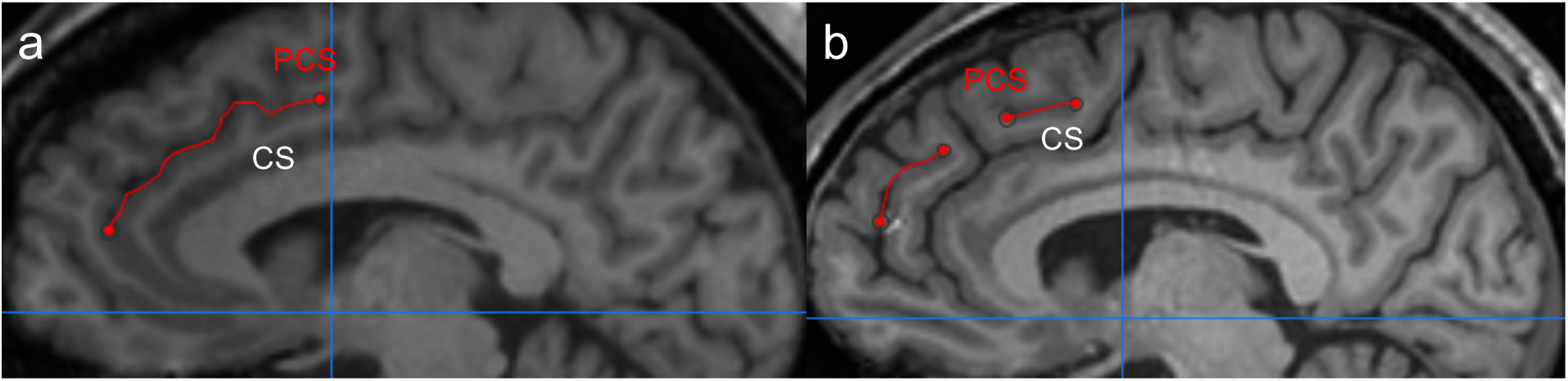
PCS measurement for two example images. Note: the paracingulate sulcus (PCS) marked in red, lies dorsal and parallel to the cingulate sulcus (CS). (a) In this image, the PCS is continuous and is measured from its origin in the first quadrant (cross-hairs at y = 0 and z = 0) to its end. (b) Here, the PCS is non-continuous, and is measured from its start in the first quadrant with inclusions of separate segments such that the total distance between them is less than 20mm.

Auditory verbal hallucinations (AVHs) are the most common modality of hallucinations reported by patients with mental health disorders, but are also experienced in the general population by around 12% of children/adolescents and 6-7% of adults (measures of life-time experience^11,12^). While AVHs appear to occur on a continuum of experience ranging from auditory imagery and intrusive thoughts to fully developed hallucinations involving the voices of other people^13^, there is evidence of both similarity and variability in their experience by individuals with and without a clinical diagnosis. For example, the phenomenology of the hallucinatory experience is generally similar in both groups in terms of localisation, loudness, and number of voices, with the perceptual experience in both cases leading to attribution of the voices to an external person or entity^14,15^. However, there are differences in frequency, duration and age of onset of hallucinatory experiences^14^, and in the preponderance of male voices^16^ and more negative content of clinical AVHs^14,17^. Indeed, a recent systematic review found that only 52% of the 21 features of hallucinations studied were experienced by individuals in both groups^18^, suggesting that the continuum model might not be an accurate representation of the variation in hallucinations experienced by individuals with and without a clinical diagnosis.

Recent research has also questioned whether healthy individuals with hallucinations show the reality monitoring impairment associated with hallucinations in patients with schizophrenia. While a meta-analysis^19^ reported differences in reality monitoring ability between hallucination-prone and non-prone healthy individuals, this analysis was based on three studies, only two of which are in the published domain^20,21^. We ourselves carried out two separate reality monitoring studies using verbal tasks, finding no evidence of a link with proneness to auditory verbal hallucinations in the general population^22^. Aynsworth et al^23^ have recently reported evidence of healthy participants with proneness to visual hallucinations showing a bias towards misremembering items that were previously presented as words as having been presented as pictures. A similar bias was also observed in patients with psychosis who experience visual hallucinations, and might indicate enhanced visual imagery in individuals prone to hallucinations.

In summary, the research linking behavioral reality monitoring impairment and hallucinations in the non-clinical population remains inconclusive. In light of the association between PCS morphology and reality monitoring ability in the healthy population^9^, and between PCS morphology and the presence of hallucinations in patients with schizophrenia^10^, we were thus motivated to investigate whether there were differences in paracingulate sulcal morphology associated with the experience of hallucinations in individuals without a clinical diagnosis. Here, we investigate PCS length in both hemispheres of the brain in three matched groups: patients with a clinical diagnosis who experience AVHs, individuals with no clinical diagnosis who experience frequent AVHs (at least once a week) but without delusions, and healthy controls, with no life-time experience of hallucinations. PCS length was measured from structural MRI scans using a previously validated measurement protocol carried out blind to group-status, with manual tracing morphological findings compared with those obtained using automated measures of local gyrification. It was hypothesized that PCS length would be shorter, and measures of gyrification around the paracingulate cortex smaller, in both clinical and non-clinical individuals with hallucinations compared with non-clinical individuals without hallucinations.

## Methods

### Participants

Fifty non-clinical participants with AVHs, 50 patients with a psychotic disorder and AVHs, and 50 healthy control subjects were matched for age, gender, handedness and years of education. There were no differences between the groups for intracranial volume (Table 1).

**Table 1.**
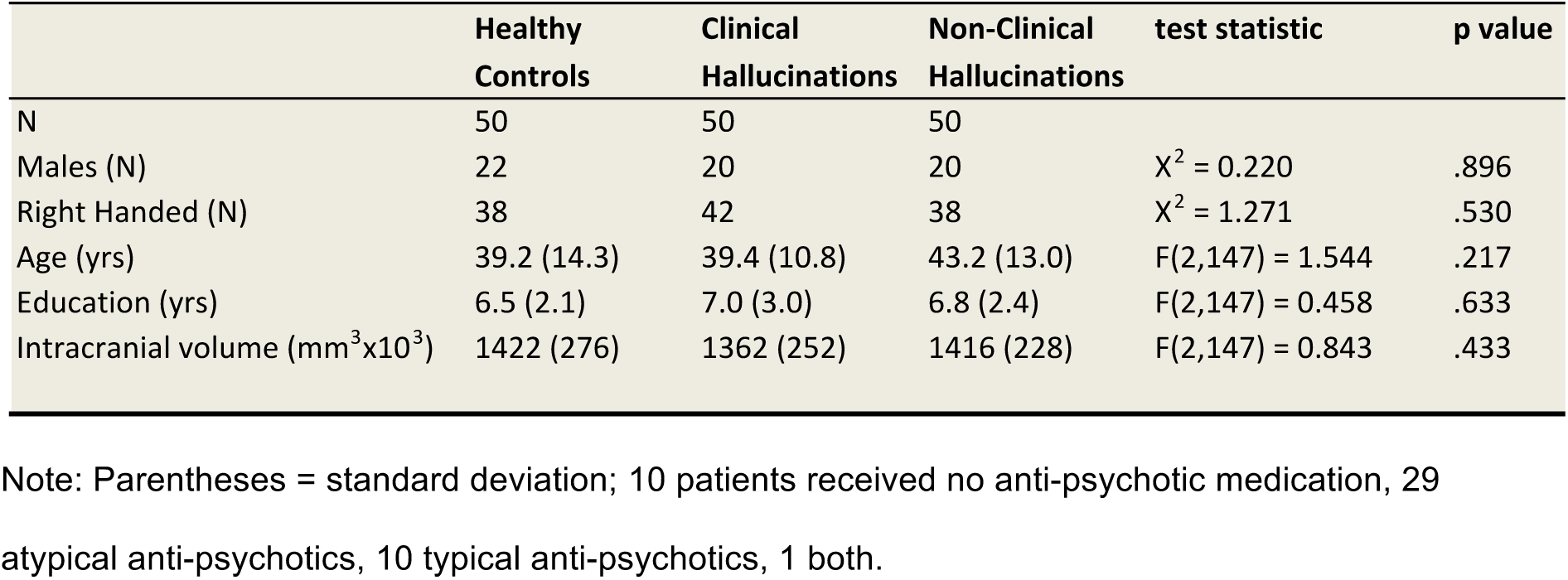
Participant data.

The recruitment procedure for healthy control and non-clinical participants with AVHs has been described previously (e.g.^24,25^). In brief, controls subjects and non-clinical individuals with AVHs were recruited via the www.verkenuwgeest.nl (‘Explore Your Mind’) website and invited for interview if they had AVH at least weekly (hallucinating group) or had not experienced AVH in their lifetime (control group). In a face-to-face interview the following inclusion criteria were checked: (i) participants had no current or past psychiatric disorders as assessed by the Comprehensive Assessment of Symptoms and History (CASH) interview and the Structured Clinical Interview for Diagnostic and Statistical Manual of Mental Disorders-III-R personality disorders (SCID-II) to exclude axis I and axis II pathology^26,27^. Importantly, hallucinating subjects were free of delusions, disorganization, and negative symptoms. Anxiety or depressive disorder in full remission were not considered exclusion criteria. Additional exclusion criteria were (ii) no chronic somatic disorder (e.g. heart failure); (iii) absence of alcohol or drug abuse for at least 3 months before the assessments. Additional inclusion criteria for participants with AVH consisted of (iv) voices that were distinct from thoughts and had a ‘hearing’ quality; (v) voices that were experienced at least once a month; and (vi) drug or alcohol abuse did not precede the first experience of AVH. All patients with a psychotic disorder were recruited from the Psychiatry Department of the University Medical Centre Utrecht, The Netherlands. Patients were diagnosed using the CASH interview according to DSM-IV criteria by an independent psychiatrist^26^. All patients met criteria for schizophrenia (29 participants), schizoaffective disorder (7), non-specific psychosis (13), or schizophreniform disorder (1), and all experienced AVHs.

Analysis of grey matter volume (GMV) differences for some of this scan data has previously been undertaken, but no analysis of paracingulate morphology has been carried out in these participants.

### Imaging data, measurement of PCS length and calculation of local gyrification index

Details of the scanning protocol, measurement of PCS length (Figure 1)^10^ and calculation of local gyrification indices^28^ are given in supplementary materials

## Results

### PCS measurement differences associated with hallucinations in clinical and non-clinical groups

There was a significant difference in total PCS length (PCS length summed across both hemispheres) between the three matched groups (patients with AVHs, non-clinical individuals with AVHs, and healthy controls), *F*(2,147) = 11.002, *p* < .001, *η*_*p*_^*2*^ *=* .130. This result survived the addition of cortical surface area for each brain scan, as a covariate, *F*(2,146) = 8.032, *p* < .001, *η*_*p*_^*2*^ *=* .099, thus controlling for a possible effect of brain size. Other potential covariates such as age, intracranial volume and global brain gyrification index had no significant effect on PCS length and were removed from the model.

Planned comparisons revealed that patients with AVHs exhibited significantly reduced PCS length compared with healthy controls, *t*(98) = 4.400, *p* < .001, *d* = .894 (mean reduction = 29.80mm), as well as with non-clinical individuals with AVHs, *t*(98) = 3.472, *p* = .001, *d* = .694 (mean reduction = 25.15mm). However, there was no significant reduction in sulcal length in non-clinical individuals with AVHs compared with healthy controls, *t*(98) = 1.013, *p* = .314, *d* = .202 (Figure 2). Consistent with earlier findings^10^, we also found main effects of hemisphere, *F*(1, 147) = 6.946, *p* = .009, *η*_*p*_^*2*^ *=* .045, but no interaction between hemisphere and group. PCS length was greater in the left hemisphere than the right hemisphere in all participant groups, t(149) = 2.647, *p* = .009, *d* = .272. Patients with AVHs exhibited shorter PCS length compared with healthy controls in both hemispheres, *t*(98) > 3.182, *p* < .002, *d* > .636, and shorter PCS length compared with non-clinical individuals with AVH, in both hemispheres, *t*(98) > 2.245, *p* < .027, *d* > .449. There was no significant difference in PCS length in either hemisphere between healthy control participants and non-clinical individuals with AVHs, *t*(98) < 303, *p* > .196, *d* < .261.

**Figure 2.**
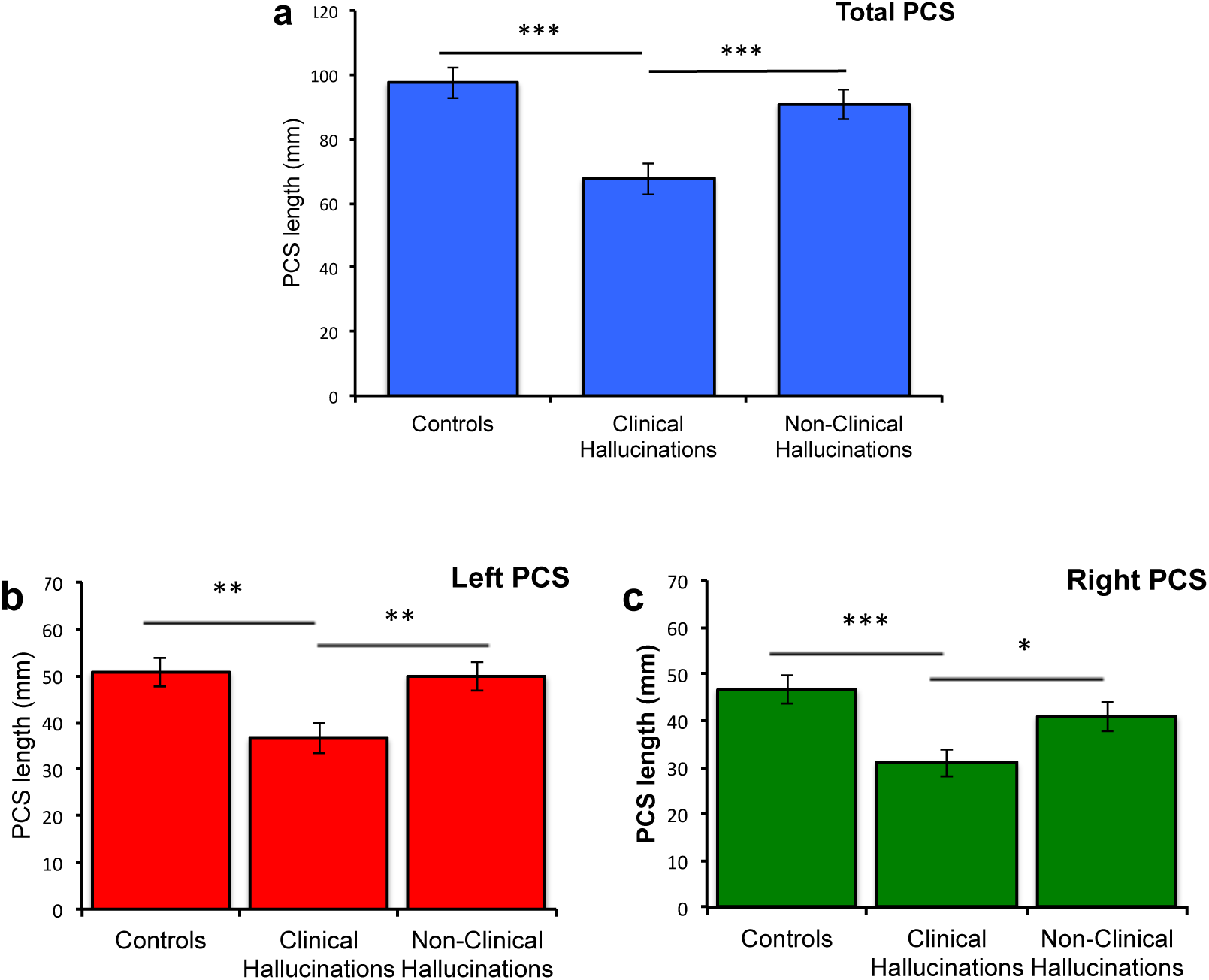
PCS length by group. (a) Total PCS length across both hemispheres (b) PCS length in the left hemisphere (c) PCS length in the right hemisphere. \*\*\**p* < 0.001, ** *p* < .0.01, * *p* < 0.05. Error bars represent standard error of the mean.

### Automated local gyrification analyses

To validate the PCS measurement findings, we conducted separate automated analyses of surface-based lGI, finding a significant reduction in mean gyrification in the mPFC regions of interest surrounding the PCS (bilateral frontopolar, medial orbitofrontal, superior frontal and paracentral cortices) in patients with AVHs when compared with healthy controls, *t*(98) = 2.128, *p* = .036, *d* = 0.425. Significant differences after correcting for multiple comparisons were also detected in the left lateral surface within the pars opercularis, inferior parietal and precentral parcellations, *t*(98) > 3.26, *p* < .001, *d* > 0.725.

We found no significant differences in mean lGI between non-clinical individuals with AVHs and either healthy controls or patients with hallucinations in these mPFC regions of interest *t*(98) < .846, *p* > .400, *d* < 0.169, (Figure 3), and no further differences across the rest of the brain that survived correction for multiple comparisons. These parcellation findings were confirmed by a whole brain analysis using a Monte Carlo procedure for multiple comparison correction. Differences in lGI were revealed in the PCS for the contrast of patients with AVHs compared to healthy controls. There were no significant clusters anywhere across the brain for the contrast of non-clinical individuals with AVHs with healthy controls, or with patients with AVHs.

**Figure 3.**
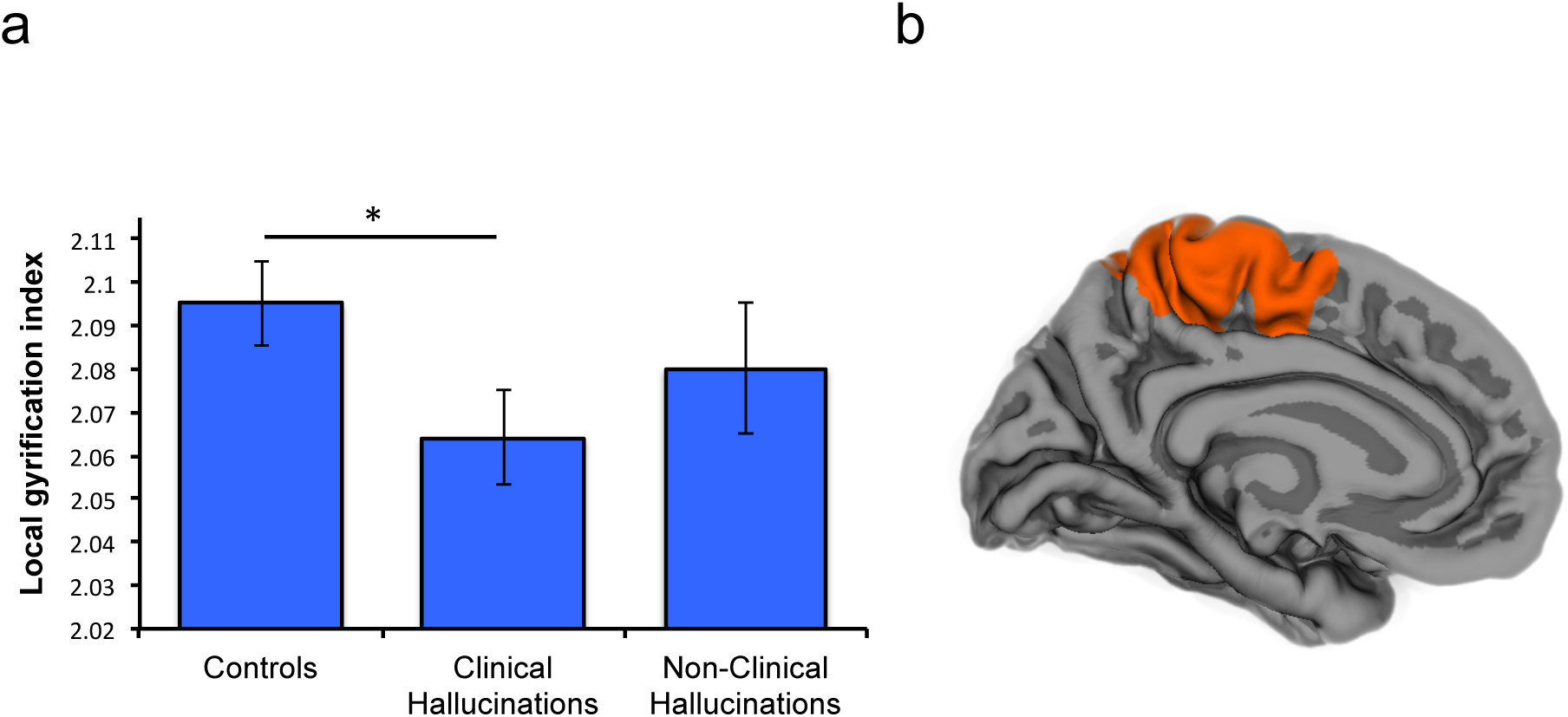
Cortical gyrification differentiates patients with AVHs, but not non-clinical individuals with hallucinations, from healthy controls. (a) Mean lGI in bilateral regions surrounding the PCS is lower in patients with hallucinations than in healthy controls, * *p* < 0.05, error bars represent standard error of the mean. (b) Local gyrification index around posterior PCS significantly differentiates clinical individuals with AVHs from healthy controls. There were no significant differences in lGI between non-clinical individuals with AVHs and either healthy controls, or patients with hallucinations.

## Discussion

In this study, we used a previously-validated visual classification technique and automated data-driven analysis to demonstrate that patients with schizophrenia who hallucinate exhibit reduced PCS length compared to both healthy controls and individuals who hallucinate in and have no clinical diagnosis. There was no difference between the hallucinating and non-hallucinating groups in our study in terms of age, sex, handedness, years of education, and brain volume. Non-clinical individuals with hallucinations had longer PCS length in both hemispheres of the brain compared with patients with hallucinations, but showed no significant difference when compared with healthy control subjects. We verified these results using a data-driven gyrification analysis, finding differences in lGI in regions surrounding the PCS between patients with hallucinations and healthy controls, but not between non-clinical individuals with hallucinations and either healthy controls or patients with hallucinations. Together, these findings suggest that the association of shorter PCS length with hallucinations is specific to patients with a psychotic disorder, with no differences in PCS length observed when comparing individuals without a clinical diagnosis who do or do not hallucinate.

These results are consistent with previous findings questioning a continuum hypothesis of hallucinatory experience, which have identified that the reality monitoring impairments associated with hallucinations in schizophrenia may not extend to non-clinical individuals with hallucinations^22^. We have suggested previously^2^ that there may be more than one route by which hallucinations occur in clinical and non-clinical groups. Hyperactivation of sensory cortices has previously been associated with the perceptual content of hallucinations (e.g. ^29,30^), and may be subject to a process of reality monitoring mediated by cortical activity within the dorsal anterior cingulate cortex (ACC) region of the mPFC. In healthy individuals without hallucinations, the sensory activity may be correctly identified by effective reality monitoring processes leading to the correct recognition of the associated perceptual content as self-generated. However, in patients with clinical hallucinations, the sensory activity may be more intense than usual^29^ (perhaps mediated by stress, trauma or tiredness^17^), and when accompanied by hypoactivation of mPFC^31^, this may then lead to reality monitoring failure to recognise the sensory activity as self-generated, resulting in the experience of a hallucination. In non-clinical individuals with hallucinations, the spontaneous activity in sensory processing areas may either be of greater intensity, or otherwise unusual, perhaps in terms of vividness, that an otherwise intact reality monitoring system might fail to recognise the stimuli as internally generated, leading to a hallucination (Figure 4.). This framework is consistent with the earlier neuroanatomical model from Allen et al^29^, but suggests a varied contribution of the different factors implicated in hallucination generation between individuals with and without a clinical diagnosis. In the case of non-clinical hallucinations, the emphasis is on the hyper-activation of sensory cortices, whereas for clinical hallucinations there is an additional effect from impairments in top-down monitoring processes.

**Figure 4.**
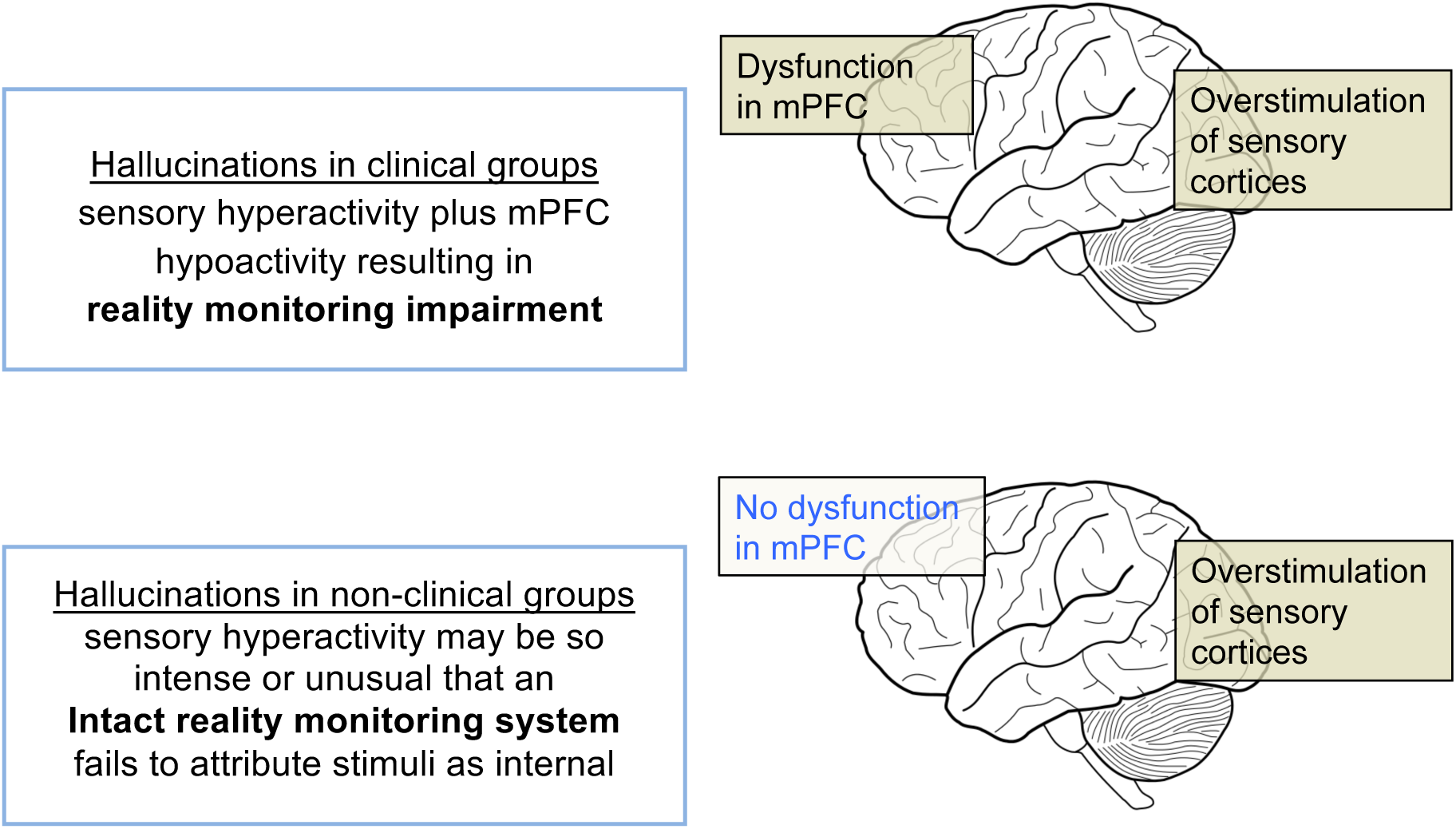
Possible mechanisms underlying hallucinations in clinical and non-clinical groups.

The suggestion of possible differential mechanisms underlying hallucinations in clinical and non-clinical groups is supported by the failure to detect dopaminergic dysfunction in non-clinical individuals with hallucinations^32^. Increased dopamine synthesis has been widely associated with the development of psychosis (e.g. ^33,34^), and suggested to relate to psychotic experience through aberrant processing of salience^35^. This failure to detect increased dopamine synthesis in non-clinical individuals with hallucinations suggests possible further differences at the neurobiological level which may be associated with the process of reality discrimination, and might relate to the phenomenological differences in the experience of hallucinations between clinical and non-clinical individuals. Investigating how changes in PCS morphology relate to dopaminergic function is an interesting future direction for this work.

Wider structural evidence also supports the association between morphological variation in the PCS and reality monitoring efficiency that underlies this model. The absence of a left hemisphere PCS in healthy individuals has been shown to be associated with a reduction in GMV in paracingulate cortex, as well as an increase in volume in the surrounding ACC^36^. Furthermore, Buda et al’s study linking impaired reality monitoring ability in healthy individuals to the bilateral absence of the PCS^9^ showed an associated GMV increase in the mPFC. These results are consistent with the observation that reduced PCS cortical folding affects both local activation patterns^37–39^, and cognitive functioning^40,41^ by altering the structural integrity of surrounding cortex. However, there may be a greater impact from reduced paracingulate folding on cognitive function due to weakened connectivity between the dorsal ACC and more distal brain regions, in particular those involved in sensory processing such as speech-sensitive auditory cortex in the superior temporal gyrus. Such differences in connectivity might arise from factors related to the process of cortical folding that occurs during gestation^42,43^. In this case, the PCS morphological differences we detect would simply be markers of the underlying cause of the associated cognitive variability. This too offers a focus for further research, with existing structural connectivity studies of hallucinations broadly supportive of such an explanation^44,45^. Furthermore, Vercammen et al^46^ found evidence of reduced functional connectivity between the left temporoparietal junction and bilateral ACC associated with more severe AVHs in patients with schizophrenia.

The proposed model of hallucinations described above is both simple and speculative, but it accounts for the broad similarity in the location of brain activity during hallucinations in clinical and non-clinical groups, particularly in the region of speech-sensitive auditory processing regions within the superior temporal gyrus^47^. The model is also consistent with failures to detect reality monitoring impairment in healthy individuals prone to hallucinations, as well as with the observation of spontaneous activity in speech sensitive auditory cortex in healthy individuals during periods of silence^48^. While imaging studies of reality monitoring typically report mPFC activity in the frontal pole, this may relate particularly to declarative task-related activity, and more posterior/dorsal ACC activity is also observed in many of these studies (e.g. ^3,4^). The proposed involvement of dorsal ACC in reality monitoring of sensory information is also consistent with a wider function of this region in error monitoring, attention, and the integration of cognitive and affective processes in executive control^49–51^.

The model can also accommodate the phenomenological differences often reported in the experience of clinical and non-clinical hallucinations. It is suggested that non-clinical hallucinations are unlikely to occur without hyper-activation of sensory cortices, which culminates in content which is unusually intense or vivid in nature. While a similar process is implicated for clinical hallucinations, this factor may be less significant, given the additional impact of impaired reality discrimination processes. Such an view can account for the lower frequency and duration of non-clinical, compared to clinical, hallucinations^14,17^, as the experience of a non-clinical hallucination might depend on the sensory information surpassing, and being maintained at a level in excess of a hallucination threshold. Furthermore, non-clinical hallucinations are associated with a greater level of control than clinical hallucinations^14,17^ which might be understandable in terms of the relative ability to down-regulate sensory activity, rather than to enhance the impaired reality discrimination process intrinsic to clinical hallucinations. It is not, however, suggested that the model can account for the entirety of the experiential differences in hallucinations between the groups; for example it cannot explain the enhanced level of negative content that may be associated with clinical hallucinations^14,15,17^.

Looking more broadly at the proposed model, there is also strong evidence supporting a role for ACC in the generation of hallucinations. Hunter et al’s^48^ demonstration of spontaneous activity in speech-sensitive auditory cortex in healthy individuals found this to also be associated with activity within the ACC. Dorsal ACC/paracingulate activity has also been related to the monitoring and generation of internal and external speech in healthy individuals and in patients with schizophrenia^37,52^. ACC activity during hallucinations has been reported in some, but not all state studies of hallucinations (e.g. ^53^), in the generation of conditioned hallucinations in healthy individuals^54^, and in self-induced hallucinations in hypnosis-prone individuals^55^. Furthermore, a recent study into the processing of ambiguous speech in individuals with non-clinical hallucinations found that voice hearers recognized the presence of speech in degraded sine-wave speech before control subjects, with the intelligibility response related to activity in both dorsal ACC and superior frontal gyrus^56^. This suggests a role for ACC in the enhanced tendency of individuals prone to hallucinations to extract meaningful linguistic content from ambiguous information.

This last finding is significant in highlighting the likely complexity of the hallucinatory process. Whilst our framework is intentionally simple, it might provide a useful basis to assist the understanding of hallucinations across clinical and non-clinical groups, and areas for future focus have been discussed above. The source monitoring framework of Johnson & Raye^1^ suggests that decisions are made as to the source of a percept through comparison of its contextual, semantic, perceptual or cognitive features with characteristic traces relating to internal or external sources^57^. However, current computational theories of hallucinations provide evidence inconsistent with the idea of a fixed percept, proposing instead a top-down driven process combining sensory information within a framework consisting of variable levels of beliefs and prior experience^58,59^. It can be argued that these are not mutually exclusive processes – while the features of a percept may be generated through a process involving top-down and bottom-up interaction, it remains a failure in source attribution which underlies the experience of a hallucination as a false percept. The relative involvement of source monitoring, attentional, and perceptual processes in the attribution of the source of sensory information remains an intriguing question, suggesting promising avenues for future research.

### Limitations

PCS length and local gyrification measures in the vicinity of the PCS for non-clinical individuals with hallucinations were intermediate between those for clinical individuals with hallucinations and control subjects without hallucinations. The failure to find a significant difference between non-clinical individuals with and without hallucinations, which would support a continuum model of hallucinations in clinical and non-clinical groups, may thus reflect an effect-size issue. Replication of this study, or use of a larger dataset would address this, but confidence on this issue is obtained from the consistency of the findings between manual measures of PCS length, and brain-wide automated gyrification analyses involving non-parametric cluster-wise correction for multiple comparisons using Monte Carlo simulation. In both cases, no significant differences were found between non-clinical individuals with and without hallucinations. In contrast, significant differences in these measures were found between clinical individuals with hallucinations and non-clinical individuals without hallucinations (above), and between patients with schizophrenia with and without hallucinations in a previous study which utilized the same techniques^10^.

In sum, we replicated earlier findings of shorter PCS in patients with hallucinations, but did not find this characteristic in non-clinical people with hallucinations. These findings, together with our previous work, corroborate of a model in which part of the mechanism underlying hallucinations is shared between clinical and non-clinical individuals with hallucinations, while a second mechanism frustrating adequate reality monitoring, is present in patients only.

