## Supplementary Materials for "Paracingulate sulcus morphology and hallucinations in clinical and non-clinical groups"

### **Imaging Data**

T1-weighted structural MRI scans were obtained using a 3T Phillips scanner, using a 3D T1 turbo field-echo sequence (repetition time 9.96ms, echo time 4.59ms, flip angle 8°, field of view 224mm, matrix 256 mm x 256 mm, voxel size .875 x .875 x 1mm). Scan time was 8 mins 50 secs.

### **Measurement of PCS length**

The PCS measurement process has been described previously^1^ and is openly available at <https://www.repository.cam.ac.uk/handle/1810/264520>). In essence, individual scans were imported as nifti folders into Mango brain visualization software (version 3.6; [http://ric.uthscsa.edu/mango)](http://ric.uthscsa.edu/mango/mango.html) and inspected for integrity. In an axial view, the locations of the anterior and posterior commissures (AC and PC) were marked, and the scan rotated to line up the AC and PC in a horizontal plane. The origin was reset to the location of the AC. On a sagittal slice, 4.375mm to the left or right of the medial line, the cingulate sulcus (CS) was identified as the first major sulcus running in an anterior–posterior direction, dorsal to the corpus callosum and typically visible for five sagittal slices or more. The PCS was then identified if present as a salient sulcus, running parallel, horizontal and dorsal to the CS, and visible for three or more sagittal slices measured from the medial plane (x = 0). The sulcus was measured using the ‘trace line’ function in Mango from its start in the first quadrant prescribed by y > 0 and the horizontal line linking the AC and PC (z > 0), starting at the point at which the sulcus ran in a posterior direction. The PCS was measured to its end point, which could fall outside of the first quadrant (Fig. 1).

#### Calculation of local gyrification index

Measures of cortical folding across the brain were obtained by calculating local gyrification indices (lGI) for each MRI structural scan using the method of Schaer et al^2^. lGI gives a measure of cortical folding by comparing the amount of cortex buried within sulcal folds at the grey/white matter interface with the amount of visible or surface cortex at each vertex of the reconstructed brain surface, based on 3D spheres of radius 25 mm. Calculations made at each vertex can then be averaged to give a global gyrification index for each hemisphere, or for 34 individual brain regions as defined by an automated parcellation procedure^3^. Calculation was undertaken using FreeSurfer Software version 5.3.0 ([http://surfer.nmr.mgh.harvard.edu](http://surfer.nmr.mgh.harvard.edu/)), which also provided measures of intracranial volume and cortical surface area for each subject’s scan, for use as covariates in the analysis. For statistical analysis, individual gyrification maps were registered to the Freesurfer average subject template. Group differences in lGI were analysed by fitting a general linear model at each vertex on the surface. Age was included as a covariate, and non-parametric cluster-wise correction for multiple comparisons was performed using Monte Carlo simulation^4^, with a threshold of p < 0.05, corrected for multiple comparisons across each hemisphere. Two-sample t-tests were used to identify region-specific differences averaged across *a priori* mPFC regions of interest in the vicinity of the PCS^3^, at a threshold of p < 0.05. Regional differences outside the region of interest were reported if they exceeded a threshold of p < 0.05, corrected for multiple comparisons across the 34 parcellated brain regions.

1.	Garrison JR, Fernyhough C, McCarthy-Jones S, Haggard M, The Australian Schizophrenia Research Bank, Simons JS. Paracingulate sulcus morphology is associated with hallucinations in the human brain. *Nat Commun*. 2015;6:8956.

2.	Schaer M, Cuadra MB, Tamarit L, Lazeyras F, Eliez S, Thiran J-P. A surface-based approach to quantify local cortical gyrification. *IEEE Trans Med Imaging*. 2008;27(2):161-170.

3.	Desikan RS, Ségonne F, Fischl B, et al. An automated labeling system for subdividing the human cerebral cortex on MRI scans into gyral based regions of interest. *Neuroimage*. 2006;31(3):968-980.

4.	Hagler DJ, Saygin AP, Sereno MI. Smoothing and cluster thresholding for cortical surface-based group analysis of fMRI data. *Neuroimage*. 2006;33(4):1093-1103.
